# Calcium-Mediated Modulation of Blood-Brain Barrier Permeability by Laser Stimulation of Endothelial-Targeted Nanoparticles

**DOI:** 10.1101/2022.06.02.494541

**Authors:** Xiaoqing Li, Qi Cai, Blake A. Wilson, Hanwen Fan, Monica Giannotta, Robert Bachoo, Zhenpeng Qin

## Abstract

The blood-brain barrier (BBB) maintains an optimal environment for brain homeostasis but excludes most therapeutics from entering the brain. Strategies that can reversibly increase BBB permeability will be essential for treating brain diseases and is the focus of significant preclinical and translational interest. Recently, we reported that picosecond laser excitation of molecular-targeted gold nanoparticles (AuNPs) induced a graded and reversible increase in BBB permeability *in vivo* (OptoBBB). Here we investigate how to increase the targeting efficiency and how picosecond lase stimulation of AuNP leads to an increase in endothelial paracellular permeability. Our results suggest that targeting brain endothelial glycoproteins leads to >20-fold higher targeting efficiency compared with tight junction targeting. We report that OptoBBB is associated with a transient elevation of Ca^2+^ that propagates to adjacent endothelial cells after laser excitation and extends the region of BBB opening. The Ca^2+^ response involves both internal Ca^2+^ depletion and Ca^2+^ influx. Furthermore, we demonstrate that the involvement of actin polymerization and Ca^2+^-dependent phosphorylation of ERK1/2 lead to cytoskeletal activation, increasing paracellular permeability. Taken together, we provide mechanistic insight into how excitation of endothelial targeted AuNPs leads to an increase in BBB permeability. These insights will be critical for guiding the future developments of this technology for brain disease treatment.

## Introduction

The vast brain neural networks and supporting glial cells are protected from circulating neurotoxins by a tightly regulated blood-brain barrier (BBB)^1^ formed by the brain capillaries. The BBB is formed by tight junctions (TJs) and adherens junctions (AJs) associated protein complexes, severely restricting paracellular diffusion^2^. That, coupled with low endothelial transcytosis^3^, results in a highly restricted movement of molecules from the circulation to the brain interstitium. While the BBB plays a protective role for the brain, it also represents a significant barrier to drug delivery. Overcoming the BBB to facilitate brain drug delivery is critical for treating central nervous system (CNS) diseases. Our recent work has demonstrated that transcranial picosecond-laser stimulation of molecular-targeted gold nanoparticles (AuNPs) can temporarily increase the BBB permeability by paracellular diffusion, referred to as OptoBBB^4^. We also showed that OptoBBB allows the delivery of various therapeutics, including human IgG, viral vector, and liposomes, into the brain *in vivo*. Thus, OptoBBB provides a promising avenue for improving brain drug delivery.

Our previous report attempted JAM-A (one of the tight junction proteins) targeting. However, the targeting efficiency is low due to JAM-A expression and limited distribution on cell boundaries. The glycoprotein is one of the critical components of endothelial glycocalyx, a dense and brush-like structure that cover the luminal surface of the BBB^5, 6^. The *Lycopersicon esculentum* lectin (LEL) can efficiently label the cerebral vessel wall by binding to glycoprotein^4, 7^. Targeting glycoprotein may be more effective for OptoBBB mediated increase in BBB permeability, further demonstrating the possibility of increasing the vascular permeability by designing new nanoparticle formulations. Previous studies showed that picosecond-laser irradiation of AuNPs causes pressure generation due to thermoelastic expansion of AuNPs^8, 9^. Tiny mechanical waves can cause G-actin polymerization into F-actin directly^10^. Mechanical forces can induce intracellular Ca^2+^ increase in endothelial cells^11, 12^. Ca^2+^, as a second messenger, plays a crucial role in the signaling pathways that regulate endothelial permeability^13^. Activating Ca^2+^-sensitive signaling has been shown to increase BBB permeability by several pathways, including ERK1/2 phosphorylation-triggered activation of actomyosin filament contraction^14^. Elucidating the contribution of Ca^2+^ and associated signaling would help clarify the mechanism for OptoBBB, which would provide additional insight into reversibly regulating vascular permeability through nanoscale mechanical and thermal perturbation.

In this study, we quantify and compare the relative effectiveness of the two targets (glycoprotein versus JAM-A) for OptoBBB. Our results suggest that glycoprotein targeting AuNPs is >20-fold more effective than tight junction targeting AuNPs. Furthermore, we test the hypothesis that Ca^2+^ signaling plays a critical role in regulating endothelial cell paracellular permeability with an *in vitro* transwell BBB model (**Figure 1a**). We show that endothelial Ca^2+^ signaling is associated with OptoBBB, and pharmacological inhibition of Ca^2+^ signaling blocks the paracellular permeability change, an accepted surrogate of the BBB regulation. Importantly, our results suggest that Ca^2+^ signals can propagate among endothelial cells and thus extend and coordinately regulate BBB opening. Further investigation of the Ca^2+^ signaling revealed the contribution of both internal Ca^2+^ release via inositol 1,4,5-trisphosphate (IP3) signaling and Ca^2+^ influx. The phosphorylation of ERK1/2 is one of the cellular events in the Ca^2+^ signaling pathway and can be detected shortly after the laser treatment (5 minutes, 0.5 hours) and recovers at 6 hours. These findings suggest that OptoBBB involves cell contraction via ERK1/2 phosphorylation through Ca^2+^ signaling and actin polymerization. This study demonstrates a method to significantly increase BBB targeting efficiency and elucidates the mechanism of OptoBBB, opening new avenues for future development of this technology and CNS disease treatment.

**Figure 1.**
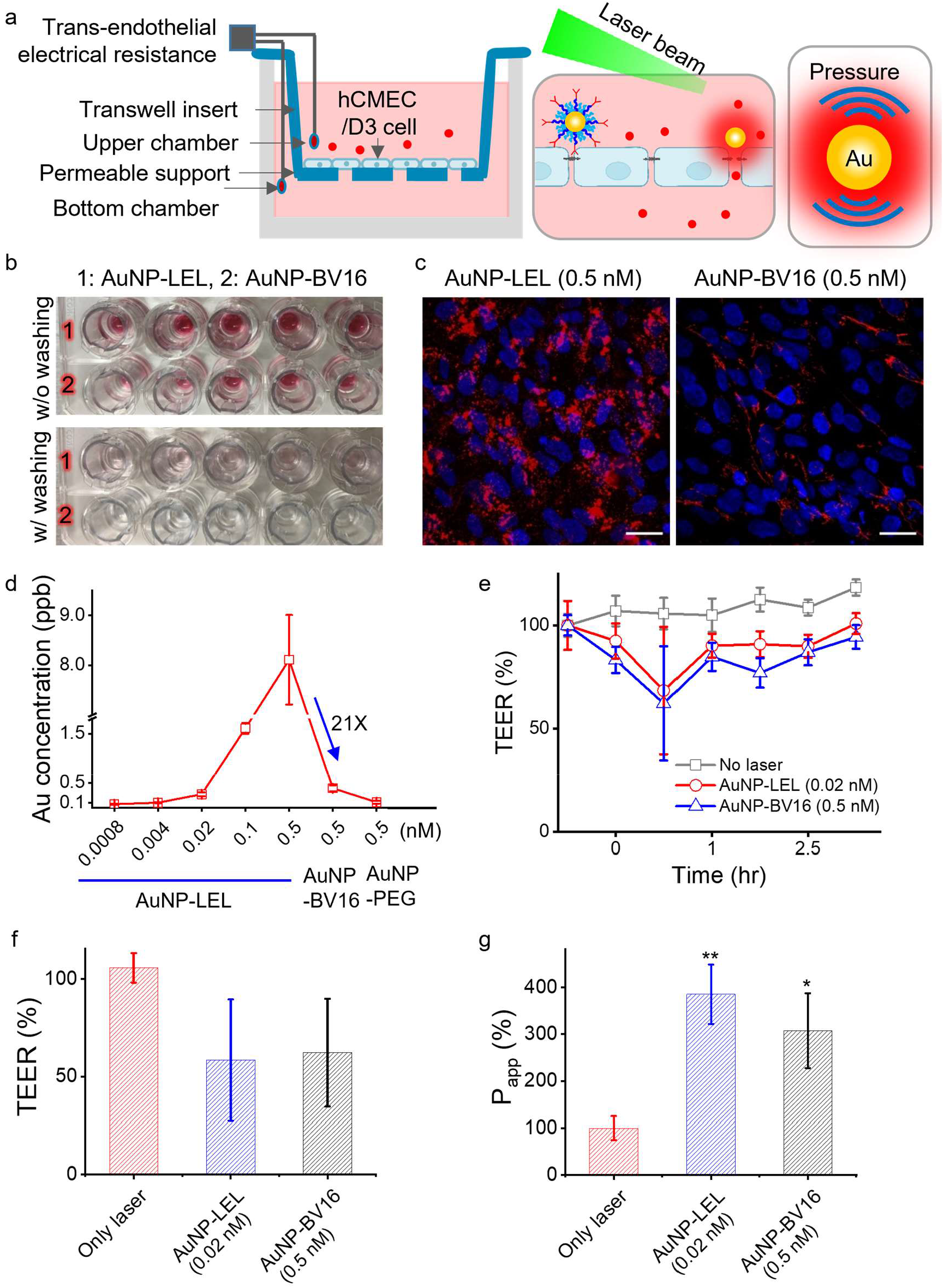
Laser stimulation of endothelial-targeted nanoparticles leads to reversible BBB permeability change (OptoBBB). **(a)** Schematic of OptoBBB *in vitro*. Red dots indicate the molecules that are impermeable to the intact BBB and become permeable to the leaking BBB after laser stimulation. **(b)** Images of the well plate after 0.5 nM AuNP-LEL and AuNP-BV16 incubation with hCMEC/D3 monolayers. **(c)** Distribution of AuNP-LEL and AuNP-BV16 (red) on hCMEC/D3 monolayers using ICC staining. Blue: nuclei. **(d)** Quantitative gold accumulation of AuNP-LEL and AuNP-BV16 on D3 monolayers using inductively coupled plasma mass spectrometry (ICP-MS), n=6. **(e)** Normalized TEER change over time by laser stimulation of AuNP-LEL (0.02 nM) and AuNP-BV16 (0.5nM), n=3. **(f)** Comparison of TEER change at 0.5 hours after laser irradiation of different targeting AuNPs, n=3. **(g)** Comparison of permeability change after laser irradiation of different targeting AuNPs, n=3. 35 mJ/cm^2^, 5 pulses, 5 Hz. Data expressed as Mean ± SD. Unpaired *t-test* was performed individually between Only laser and the other groups. *: P<0.05, or **: P<0.01, was considered a statistically significant difference. Scale bar: 20 μm.

## Results

### Glycoprotein targeting leads to significantly improved AuNP accumulation on BBB

First, we tested the hypothesis that targeting glycoprotein would lead to a more effective targeting efficiency to endothelial cells due to its abundant expression over the luminal surface of the microvasculature relative to JAM-A, which is restricted to the tight-junction protein complex. We adapted a transwell model using human cerebral microvascular endothelial (hCMEC) D3 cell^15, 16^ for our study. Its associated trans-endothelial electrical resistance (TEER), permeability, and TJ formation have been extensively validated (**Figure S1**), consistent with the previous report^17^. We prepared the tight junction targeting AuNPs with anti-JAM-A antibody (AuNP-BV16) and glycoprotein-targeting AuNPs with lectin (AuNP-LEL) (**Figure S2**) using polyethylene glycol (PEG) backfilling^18^. When incubating AuNP-BV16 and AuNP-LEL with D3 monolayers using the same concentration (0.5 nM), a significantly higher amount of AuNP-LEL binds to the cell surface than AuNP-BV16, as indicated by the pink color differences after washing with PBS to remove free AuNPs (**Figure 1b)**. The ICC staining confirmed the higher accumulation of AuNP-LEL (**Figure 1c**). We further measurement gold concentration and found that AuNP-LEL (0.5 nM) displays 21 times higher accumulation on the D3 cells than AuNP-BV16 at the same concentration (**Figure 1d**). We then selected a much lower AuNP-LEL dose (0.02 nM) for the following experiments as it shows a similar gold accumulation to 0.5 nM AuNP-BV16.

We then compared the BBB opening with AuNP-BV16 and AuNP-LEL and laser excitation. We tested different conditions for BBB opening *in vitro*. We selected the optimized combination (35 mJ/cm^2^, 5 pulses, 5 Hz, and 50 nm AuNPs) for the following experiments to achieve a high BBB opening efficiency (indicated by more significant TEER drop and higher permeability) with minimal cell injury (**Figure S3–S7**). OptoBBB with 0.02 nM AuNP-LEL and 0.5 nM AuNP-BV16 shows comparable TEER and permeability changes (**Figure 1e–g)** without causing significant cell injury **(Figure S8**). Furthermore, we tested the BBB modulation *in vivo* with these two targets. The gold element analysis shows that AuNP-LEL accumulated 12-fold higher in the brain than TJ-targeting AuNPs (AuNP-BV11) (**Figure S9a**). Under the same AuNP dose and same laser pulse energy, AuNP-LEL leads to a more efficient BBB opening, indicated by a larger area of Evans blue leakage (**Figure S9b**). Thus, targeting glycoproteins increases AuNP accumulation on the BBB and allows a lower concentration of AuNPs to reach comparable efficacy compared with TJ-targeting AuNPs.

### Laser-induced elevation of Ca^2+^ propagates among endothelial cells and leads to BBB opening

Second, we examined the Ca^2+^ signaling and propagation during OptoBBB. We imaged intracellular free Ca^2+^ signals using fluo-4 as an indicator before and after a focused laser stimulation (**Figure 2a**). D3 monolayers were incubated with endothelial-targeting AuNPs and fluo-4 before imaging. After washing with PBS, we place the monolayers in the relevant medium under a real-time fluorescent imaging system. We then imaged 30-50 seconds before laser stimulation as a baseline and 120-180 seconds after laser excitation. Our results show Ca^2+^ level started to increase at 2 seconds after laser excitation (**Figure 2b, c, S10, and video S1**). In contrast, no laser stimulation shows no increase in fluo-4 intensity over the baseline. A similar result was seen with only laser excitation in the absence of AuNPs (**Figure 2c**). Interestingly, the image analysis shows Ca^2+^ propagation from laser-irradiated regions to adjacent and then distant regions (**Figure 2b,d**, **S10c,d**, and **video S1**). We further observed that the increased Ca^2+^ signal in some cells oscillated (**Figure S10e**,**f**), potentially resulting from a certain IP3 (an agonist to trigger Ca^2+^ release from endoplasmic reticulum) concentration to support this oscillatory event^19^. Importantly, incubation of the monolayers with BAPTA (Ca^2+^ chelator, 10 μM) blocks Ca^2+^ signaling and completely abolishes the TEER changes (**Figure 2e**). These findings confirm the involvement of transient Ca^2+^ elevation in the reversible BBB opening, and that blocking Ca^2+^ is sufficient to block the BBB opening.

**Figure 2.**
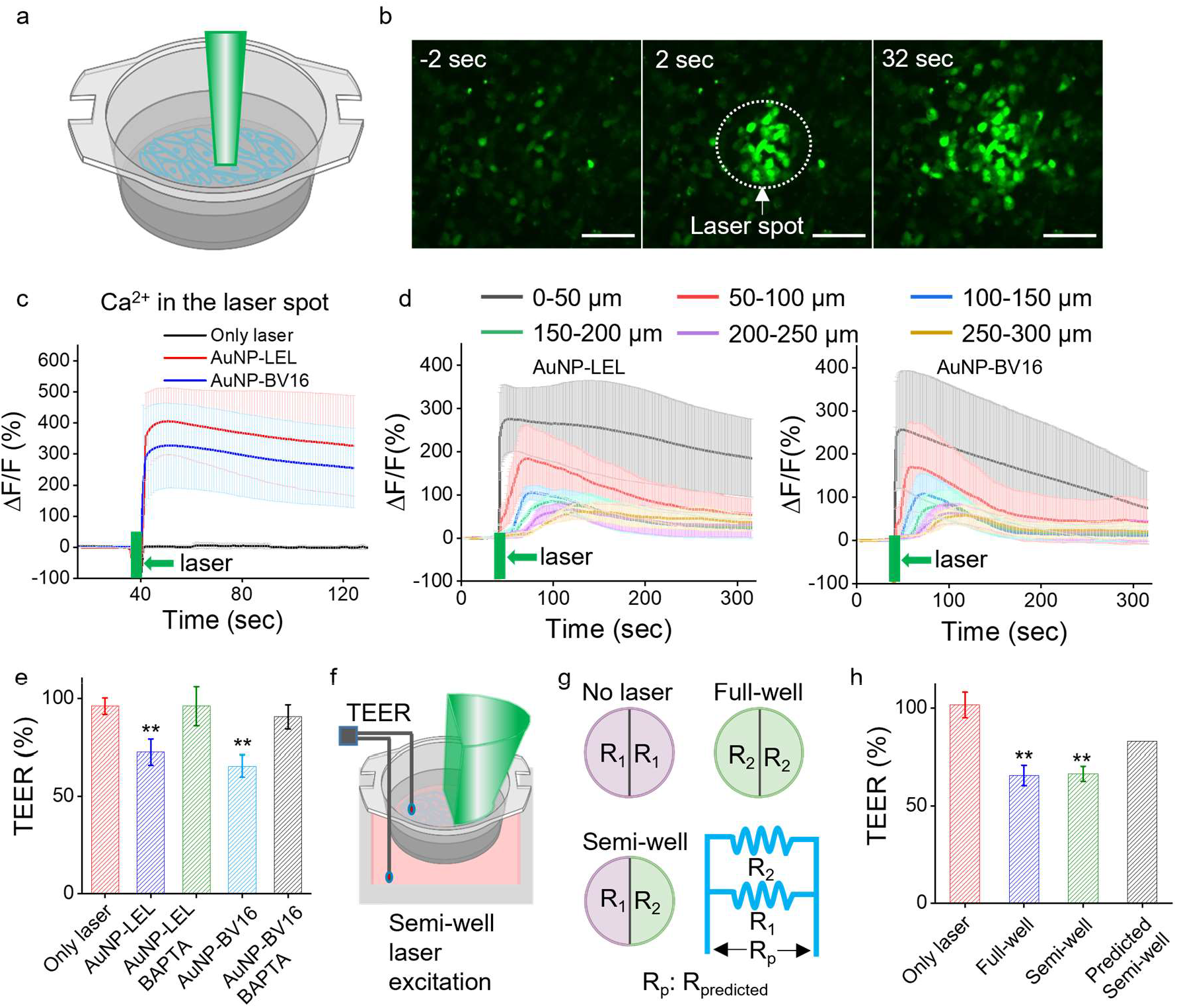
Ca^2+^ elevation and propagation among endothelial cells. **(a)** Schematic of Ca^2+^ imaging with a focused laser beam. **(b)** Representative images of the fluorescent intensity of fluo-4 (Ca^2+^ indicator) before and after laser stimulation of AuNP-LEL. Laser stimulation time is defined as 0 seconds. Green: fluo-4. **(c)** Transient Ca^2+^ elevation in the laser spot after laser stimulation, 35 mJ/cm^2^, 1 pulse. **(d)** Analysis of Ca^2+^ signaling shows the propagation after laser stimulation of AuNP-LEL and AuNP-BV16. 35 mJ/cm^2^, 1 pulse. 30-60 cells were analyzed from 3 experiments. **(e)** Ca^2+^ chelator (BAPTA) prevents TEER drop. 35 mJ/cm^2^, 10 pulses **(f)** The schematic shows only the right semi-well receives light. **(g)** A parallel circuit model for predicting the resistance of semi-well excitation is no propagation case. **(h)** Comparison of TEER with semi-and full-well laser stimulation and predicted resistance. 35 mJ/cm^2^, 10 pulses. Ca^2+^ data corresponding to 30-60 cells was analyzed from 3 experiments. TEER data correspond to 3 replicates. Data expressed as Mean ± SD. Unpaired *t-test* was performed individually between Only laser and the other groups. **: P<0.01was considered a statistically significant difference. Scale bar: 100 μm.

Ca^2+^ propagation has been reported in endothelial cells^20, 21^ and can amplify the BBB opening^22^. To investigate the contribution of Ca^2+^ propagation on OptoBBB, we examined the TEER changes by blocking the laser irradiation on half of the transwell (**Figure 2f**). A parallel circuit model (**Figure 2g**) predicts the resistance change by calculating the parallel resistance across the two semi-wells. The experimental result shows that semi-well laser excitation leads to a similar TEER drop with the full well excitation. However, the predicted resistance change of semi-well excitation is much less than the experimental measurement (**Figure 2h**). This finding indicates that Ca^2+^ propagation extends the area of BBB opening. Taken together, our results suggest that the elevation and propagation of cytosolic Ca^2+^ contribute to OptoBBB.

### OptoBBB involves both internal Ca^2+^ depletion via IP3 signaling and extracellular Ca^2+^ influx

Next, we investigated the role of internal Ca^2+^ depletion and extracellular Ca^2+^ influx during OptoBBB, as both can lead to Ca^2+^ signaling for endothelial cells (**Figure 3a**). To study the contribution of intracellular Ca^2+^ release, we removed the extracellular Ca^2+^ by incubating the cells with a Ca^2+^-free medium. Ca^2+^ elevation persists for both AuNP-LEL and AuNP-BV16 cases (**Figure 3b**). Previous studies reported that cell membrane deformation by mechanical stimulation triggered an increase in cytosolic IP3 that binds to inositol trisphosphate receptor (IP3R) on the endoplasmic reticulum (ER) to trigger Ca^2+^ release from the ER pool^23-26^. To further confirm the role of IP3 signaling, we pretreated the monolayers using 2-aminoethoxydiphenyl borate (2-APB: 200 μM), an IP3R blocker and observed no Ca^2+^elevation after laser irradiation in Ca^2+^-free medium (**Figure 3b**). As a control experiment and previously discussed, pretreatment of the cells with BAPTA, a high-affinity Ca^2+^ chelator, ablates the Ca^2+^ response (**Figure 3b**).

**Figure 3.**
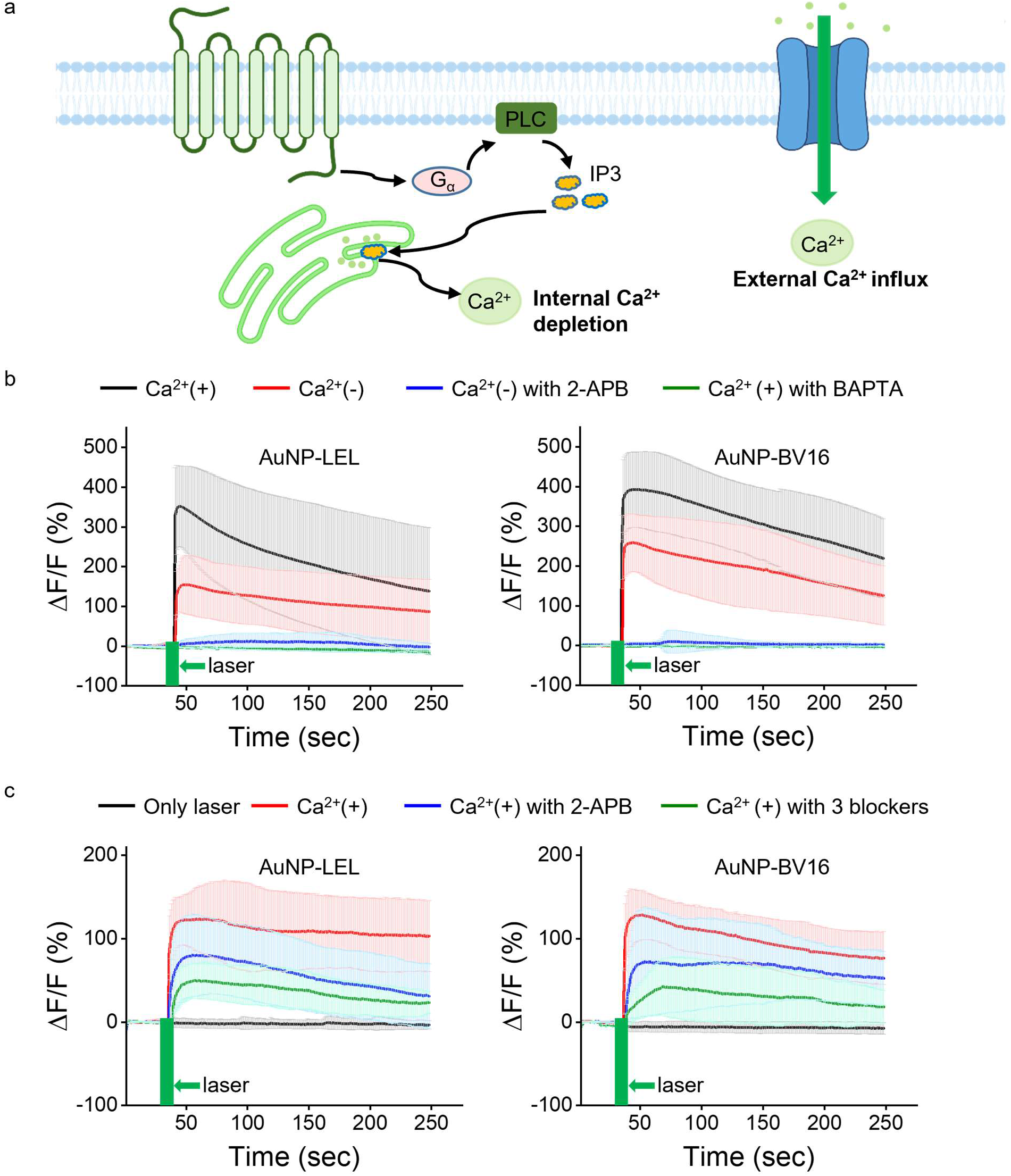
OptoBBB involves internal Ca^2+^ depletion and Ca^2+^ influx. **(a)** The schematic of internal Ca^2+^ depletion via IP3 signaling and Ca^2+^ influx. **(b)** Internal Ca^2+^ depletion after laser stimulation of AuNP-LEL and AuNP-BV16. 35 mJ/cm^2^, 1 pulse. Ca^2+^(+): FBS-free D3 culture medium that contains Ca^2+^ ions. Ca^2+^(-): HBSS solution without Ca^2+^ ions to remove the Ca^2+^ source from the extracellular environment. Ca^2+^(-) with 2-APB: HBSS solution without Ca^2+^ ions and pretreatment with IP3R blocker (2-APB) to suppress internal Ca^2+^ release. Ca^2+^(+) with BAPTA: FBS-free D3 culture medium and pretreatment with BAPTA (Ca^2+^ chelator). **(c)** Ca^2+^ influx after laser stimulation of AuNP-LEL and AuNP-BV16. Ca^2+^(+) with 2-APB: FBS-free D3 culture medium and pretreatment with IP3R blocker. Ca^2+^(+) with 3 blockers: FBS-free D3 culture medium and pretreatment with 2-APB, GSK2193874, and GsMTx4 to block IP3 signaling, TRPV4, and Piezo 1, respectively. 30-60 cells were analyzed from 3 experiments. Data expressed as Mean ± SD.

We further studied the contribution of extracellular Ca^2+^ influx on the cytosolic Ca^2+^elevation. We blocked the internal Ca^2+^ source using 2-APB and observed a transient Ca^2+^ increase in the Ca^2+^-containing medium. We observed no Ca^2+^elevation when switched to a Ca^2+^-free medium. This result suggests that Ca^2+^ influx from the extracellular solution (**Figure 3c**) contributes to the observed cytosolic Ca^2+^ response. TRPV4 and Piezo 1 are two mechano-sensors at the cell membrane to sense the mechanical stimuli and trigger many cellular processes, including Ca^2+^ influx. To further investigate the contributions of ion channels for Ca^2+^ influx, we pretreated monolayers with TRPV4 and Piezo 1 inhibitor (GSK2193874: 10 μM, GsMTx4: 10 μM) in the presence of the IP3R blocker, and the Ca^2+^ elevation persisted although with a smaller magnitude. This result indicates that other ion channels in addition to TRPV4 and Piezo 1 on the plasma membrane or changes in the plasma membrane itself may contribute to the observed Ca^2+^ influx. These studies suggest that OptoBBB involves internal Ca^2+^ depletion through IP3 signaling and extracellular Ca^2+^ influx through multiple mechanisms.

### OptoBBB involves ERK1/2 phosphorylation and actin polymerization

Lastly, we explored the downstream signaling associated with increased BBB permeability, including actin polymerization and ERK1/2 phosphorylation^27-29^(**Figure 4a**). Western blotting results show increased p-ERK1/2 signal following laser stimulation with onset at 5 minutes, peak at 0.5 hours, and a return to baseline at 6 hours (**Figure 4b,c**). Application of p-ERK1/2 blocker U0126 suppressed the p-ERK1/2 level (**Figure 4b,c**), consistent with previous reports^30^. Rapid endothelial cytoskeletal rearrangement facilitates BBB disruption^29^. To assess the actin skeleton response, the F-actin was stained by phalloidin after laser excitation of AuNPs. The results show that the F-actin fiber signal increases at 5 minutes, maximizes at 0.5 hours, and returns to baseline at 6 hours (**Figure 4d and S11**). Quantitative analysis shows an increase in F-actin immunofluorescence and a decrease in the anisotropy score of F-actin, at 5 minutes and 0.5 hours, and recovery at 6 hours (**Figure 4e,f**). The increased immunofluorescence indicates more F-actin formulation after laser stimulation. The decreased anisotropy score suggests F-actin fibers reorientate from more directions. A strongly correlated temporal profile of p-ERK1/2 signaling and F-actin polymerization does not suggest a direct causal link. These findings suggest OptoBBB involves phosphorylation of ERK1/2 and actin polymerization.

**Figure 4.**
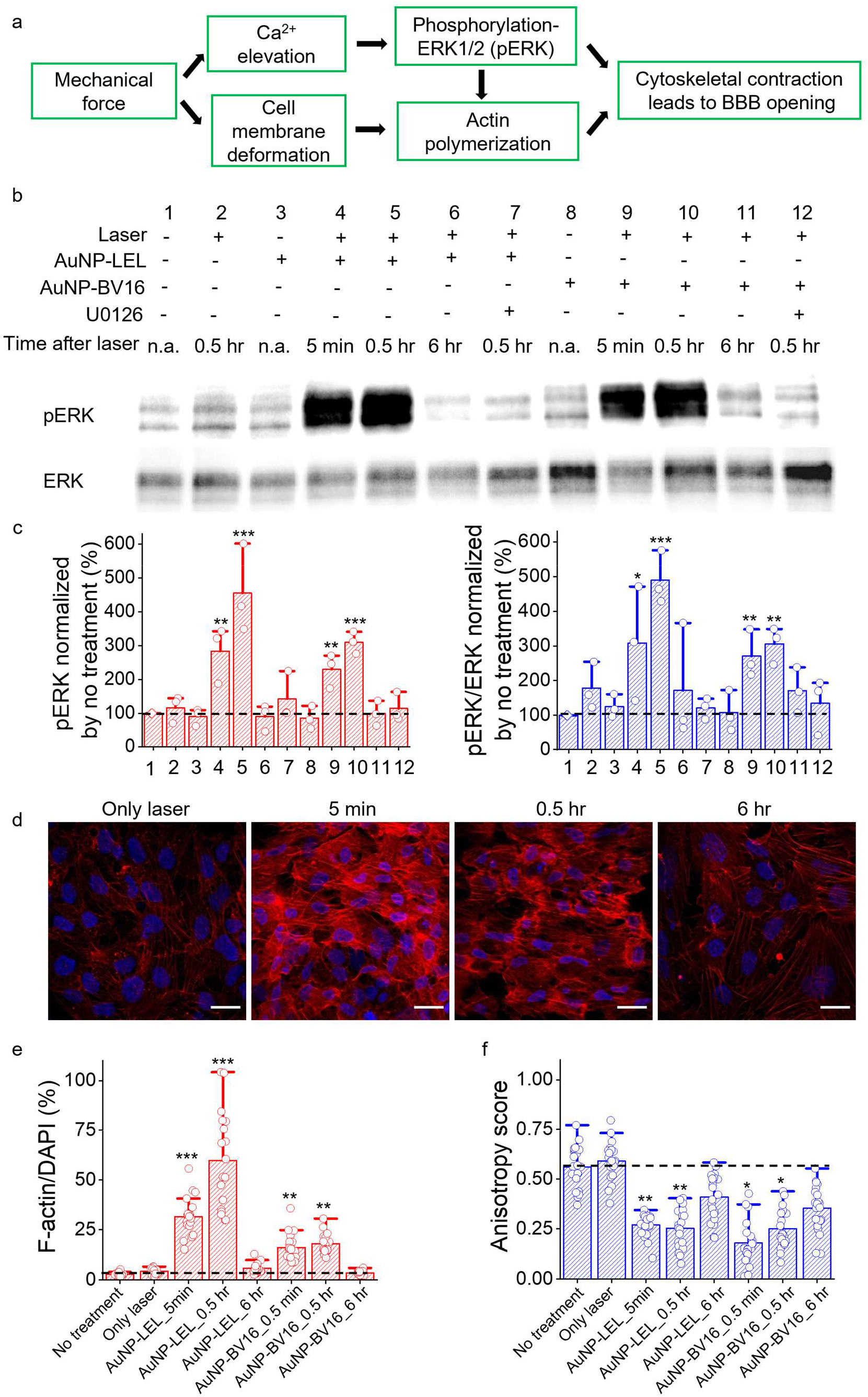
OptoBBB involves ERK1/2 phosphorylation and action polymerization. **(a)** Proposed signaling pathway for OptoBBB. **(b)** Phosphorylation of ERK1/2 (pERK) after laser excitation detected by western blot. 35 mJ/cm^2^, 1 pulse. **(c)** Quantification analysis of data in (b). Left panel: pERK normalized by control, No treatment. Right panel: ratio of pERK and total ERK1/2 (ERK) normalized by control, No treatment. **(d)** F-actin staining by phalloidin after laser stimulation of AuNP-LEL. 35 mJ/cm^2^, 1 pulse. **(e)** Quantification of fluorescent intensity of F-actin normalized by nuclei. Red: F-actin. Blue: nuclei. **(f)** Quantification of anisotropy score of F-actin. The black dash line indicates the baseline. Western blot data correspond to 3 replicates. F-ctin data correspond to 21-30 field of views (FOVs) from 3 experiments. Data expressed as Mean ± SD. Unpaired *t-test* was performed between No treatment and the other group, respectively. *: P<0.05, **: P<0.01, or ***: P<0.001 was considered a statistically significant difference. Scale bar: 20 μm.

## Discussion

Our results lead us to hypothesize the following mechanistic framework for OptoBBB (**Figure 5**). Pulsed laser excitation of vascular-targeting AuNPs produced nanoscale mechanical and thermal perturbation. This tiny perturbation triggers (1) actin polymerization; (2) Ca^2+^-influx; (3) mechano-sensor GPCR activation resulting in G protein activity, which in turn increases the production of cytosolic IP3. IP3 activates the receptors on the endoplasmic reticulum, leading to the Ca^2+^ release from the ER. The elevation of Ca^2+^ from the influx and IP3 pathway activates ERK1/2 phosphorylation. The phosphorylation of ERK1/2, together with the actin network, leads to an activation of the cytoskeleton resulting in an increase in paracellular permeability.

**Figure 5.**
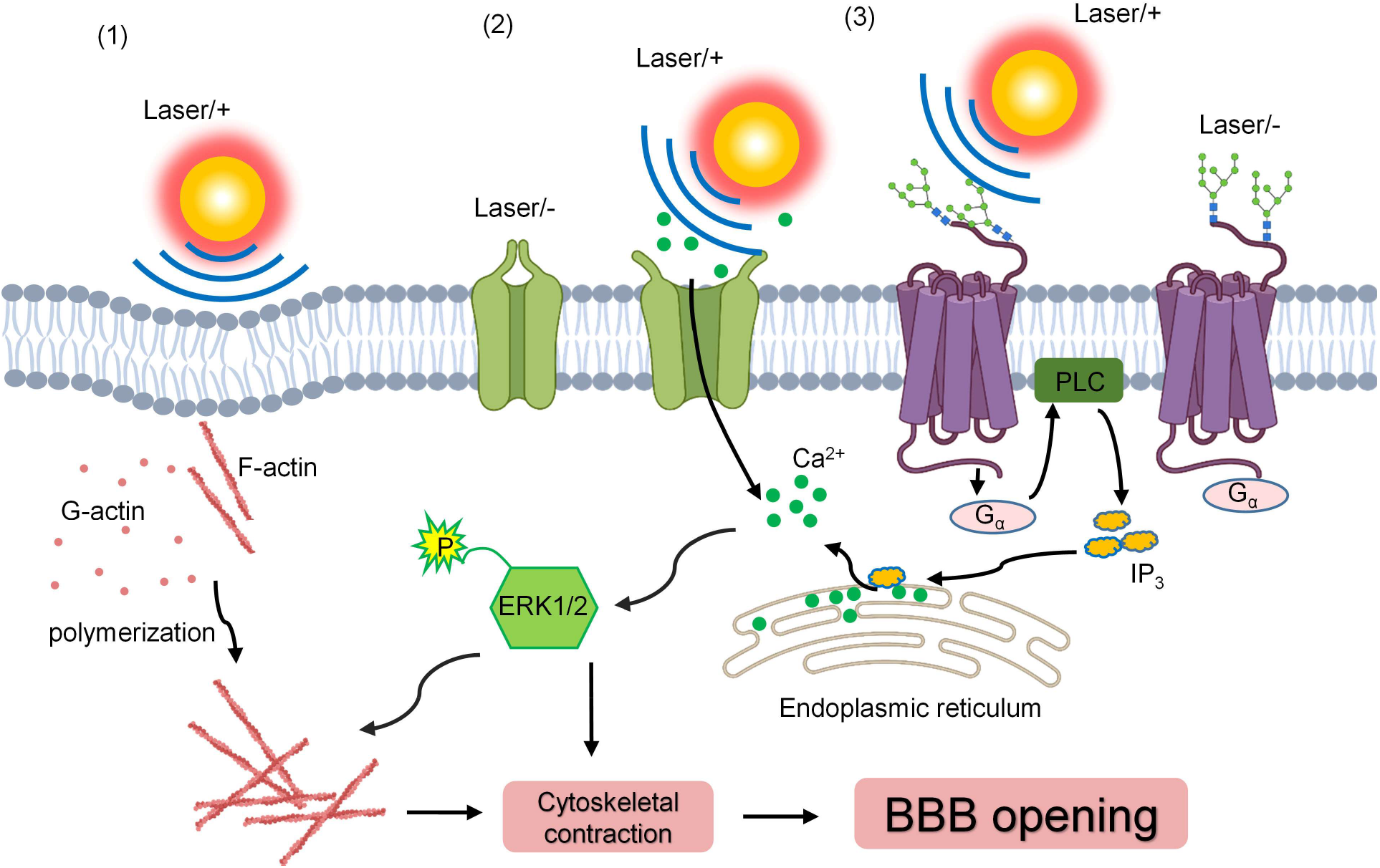
Summary of the proposed mechanism for OptoBBB. The mechanical pressure generated by laser excitation of AuNPs produces 3 effects: (1) The mechanical pressure gently deforming the cell membrane triggering actin polymerization, (2) Ca^2+^-influx from mechanosensitive channels, and (3) mechanosensitive GPCR activation resulting in G protein activity, which in turn increases the production of cytosolic IP3. IP3 activates the receptors on the endoplasmic reticulum (ER), leading to the Ca^2+^ release from the ER. Then the elevation of Ca^2+^ activates the downstream ERK1/2 phosphorylation. Finally, the phosphorylation of ERK1/2 together with actin networks causes a cytoskeletal contraction to increase the paracellular space between adjacent cells, eventually increasing the BBB permeability. Laser/-: only targeting AuNPs without laser stimulation. Laser/+: targeting AuNPs stimulated by picosecond laser.

Our *in vivo* investigation showed that the increased BBB permeability involves paracellular diffusion through the TJs, indicated by the fact that lanthanum nitrate (electron enhance tracer) filled the TJ clefts and diffused into the basement membrane and brain extracellular space^4^. In this work, the findings show a drop in TEER after laser stimulation of endothelial-targeting AuNPs. The TEER drop suggests solutes transporting from the upper chamber to the bottom chamber through paracellular space, consistent with the *in vivo* observation. However, we cannot rule out that transcytosis contributes to the increased BBB permeability. A more comprehensive evaluation of transcytosis could be carried out by analyzing the cellular uptake of fluorescent molecules (FITC-dextran) using confocal imaging and flow cytometry^31, 32^.

Glycoprotein is one of the molecular components of the glycocalyx, a thick matrix highly expressed on the luminal surface of the BBB^33, 34^. Targeting glycoproteins is expected to increase the targeting efficiency since glycoproteins display a much larger surface area on the BBB than the tight junction. Our results show significantly higher AuNPs accumulation for AuNP-LEL than AuNP-BV16 *in vitro* (21-fold) and *in vivo* (12-fold). We observed that targeting glycoprotein leads to more efficient BBB opening than targeting JAM-A with the same AuNP dose. Future studies may further investigate intracarotid infusion of AuNP-LEL to increase the targeting efficiency of AuNP-LEL *in vivo*. The high targeting efficiency enables a much lower AuNP dose required for BBB opening (0.02 nM AuNP-LEL versus 0.5 nM AuNP-BV16) and higher BBB opening efficiency.

## Conclusion

We studied the BBB targeting and cellular mechanism of OptoBBB. Specifically, targeting the glycoprotein significantly improves the vascular targeting efficiency compared with targeting tight junction *in vitro* and in *vivo*. We further observed the elevation and propagation of Ca^2+^ among endothelial cells and actin polymerization after laser excitation of endothelial-targeting AuNPs. We detected phosphorylation of ERK1/2 on the Ca^2+^-dependent signaling pathway. The Ca^2+^-sensitive signaling and actin polymerization lead to cytoskeletal activation and increase the paracellular space and permeability. Our study demonstrates a method to significantly increase BBB targeting efficiency and elucidates the calcium-dependent mechanism for OptoBBB, opening new avenues for future development of this technology and CNS disease treatment.

## Materials and Methods

### Materials

Anti-JAM-A antibodies BV16 and BV11 were provided by Dr. Monica Giannotta at the FIRC Institute of Molecular Oncology Foundation. Human cerebral microvessel endothelial cell/D3 cell line, EndoGRO™-MV Complete Media Kit, FGF-2, trypsin-EDTA, collagen type I, and millicell ERS-2 System were purchased from Millipore. LF PVDF transfer kit stain-free protein gel and western ECL substrate were purchased from Bio-Rad. P-p44/42 ERK1/2 and p44/42 ERK1/2 were purchased from Cell Signaling Technology. Gold (III) chloride, FITC-dextran, DMSO, hydroquinone, sodium citrate tribasic, BSA, Tween 20, Triton-X 100, sodium carbonate, sodium bicarbonate, phalloidin-RF, and sucrose were purchased from Sigma-Aldrich. Penicillin-streptomycin and donkey anti-mouse IgG (H+L) 488 were purchased from Life Technologies. RIPA buffer, protease inhibitor cocktail, phosphatase inhibitor cocktail, and BCA protein assay kit, fluo-4 AM, OPSS-PEG-SVA, and mPEG-SH were purchased from Laysan Bio, Inc. Biotin-PEG-SH was purchased from NANOCS. Cy3-labeled streptavidin, peroxidase-conjugated affiniPure goat anti-rabbit IgG (H+L), donkey serum, goat serum, Trypan blue, gold reference standard solution, WST-1 kit, U0126, GSK2193874, GsMTx4, BAPTA-AM, fluo-4, Hoechst dye 33342, Dulbecco’s phosphate-buffered saline, 20 kDa dialysis membrane, 6-, 24-, 96-well plates, and transwell inserts were purchased from Thermo Fisher Scientific. All other chemicals were analytical grade. Adult mice were ordered from Charles River Laboratories. Animal protocols were approved by the Institutional Animal Care Use Committee (IACUC) of the University of Texas at Dallas.

### Formation and characterization of cellular monolayers

Human cerebral microvessel endothelial cell/D3 (hCMEC/D3) cells were cultured in EndoGRO™-MV Complete Media supplemented with FGF-2 on a collagen-coated porous membrane. Once plated to the transwell insert of the 24-well plate and 96-well plate, the D3 cells were cultivated for approximately 6 to 7 days to form cellular monolayers at 37 °C and 5% CO_2_. The hCMEC/D3 monolayers were characterized by trans-endothelial electrical resistance (TEER), permeability, and immunocytochemistry (ICC) staining.

### Transendothelial electrical resistance (TEER) measurement

The electrical resistance of hCMEC/D3 monolayers was measured in Ohms (Ω). A cellular monolayer was formed on transwell inserts (0.3 cm^2^, pore size 8 μm), including upper and bottom compartments. For TEER measurement, cellular monolayers were detected by an epithelial voltmeter (Millicell ERS-2, Millipore, USA). Rinsing of electrodes by cell culture medium was required between blank-well and sample-well. The TEER values of cellular monolayers were measured before laser (−0.5 hours) and at various time delays after laser irradiation (0, 0.5, 1, 1.5, 3, and 6 hours). The resistance value of blank culture inserts coated with collagen on the top side of the membrane was used as the baseline. To obtain TEER, the baseline value was subtracted from the resistance measured from the cell monolayer samples. The resulting resistance value multiplied by the effective membrane area gives the TEER value in Ω·cm^2^.

### Permeability measurement

All the medium of cellular monolayers was replaced by a cell culture medium without 5% FBS from the top and chambers, followed by 30 minutes of incubation in 37 °C and 5% CO_2_ incubator. After that, a medium mixed with 0.3 mL FITC-dextran (40 kDa, 1 mg/mL) was added to the upper compartments. At particular time points, 100 μL of the medium was aspirated from the bottom well and added to 96 black well plates. 100 μL of culture medium was then replaced in the bottom chamber to keep the volume constant. We then measured the fluorescent intensity for the collected samples in the 96-well plate (excitation at 490 nm and emission at 540 nm) to obtain FITC-dextran concentration. The quantity (Q) was obtained by multiplying the concentration and volume. The apparent permeability (P_app_, cm/sec) was calculated by the rate of FITC-dextran quantity change over time (dQ/dt), divided by the initial concentration of dextran (C) and the membrane area (A).

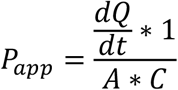

### Immunocytochemistry (ICC) staining

The JAM-A expression on hCMEC/D3 cells was characterized by ICC staining. Cellular monolayers were fixed for 5 minutes in pure methanol on ice and then washed in PBS 3 times on a shaker. A blocking buffer (5% donkey serum 2% BSA in PBS 0.05% Tween) was applied at room temperature (RT) for 1 hour. Then the cells were incubated with primary antibodies, mouse anti-human JAM-A (BV16: 3-5 μg/mL), overnight at 4°C or room temperature for 1 hour, followed by a second antibody incubation room temperature for 1 hour. Finally, the samples were incubated with Hoechst dye for 10 minutes in the dark to stain the nuclei. The samples were washed in PBS every time before changing reagents.

For F-actin staining, all samples were fixed with 4% paraformaldehyde (PFA) for 10 minutes at 4°C. After washing with PBS, treated monolayers were incubated with phalloidin-RFP (diluted ratio: 1:1000) at RT for 1 hour, followed by incubation of Hoechst dye. The monolayers were then mounted on glass slides after washing.

To compare the distribution of AuNP-BV16 and AuNP-LEL on hCMEC/D3 cells, AuNP-BV16 and AuNP-LEL were backfilled by biotin-PEG-SH (PG2-BNTH-1K, NANOCS) to stabilize the nanoparticles. AuNP-BV16-biotin and AuNP-LEL-biotin were incubated with monolayers for 0.5 hours at 37°C. After washing with PBS 3 times, the treated monolayers were fixed with 4% PFA for 10 minutes at 4°C. Cy3-labeled streptavidin (1:200) was used to detect biotin and thus the distribution of targeting AuNPs. Confocal microscopy (FV3000RS or SD-OSR) was utilized to take fluorescent images after ICC staining.

### Gold nanoparticle conjugation

AuNPs were synthesized following a previously reported method^11^. BV16, an anti-JAM-A antibody, was diluted to 0.5 mg/mL in PBS, followed by dilution in aqueous 10mM NaHCO_3_ at pH8.5. OPSS-PEG-NHS was dissolved in NaHCO_3_ and quickly added to the diluted antibody at a 125:1 molar ratio. The mixture was vortexed briefly and kept for 3 seconds, shaking on ice, followed by dialysis for 3 hours to remove free OPSS-PEG-NHS through a 20 kDa MW membrane. The thiolated BV16 was reacted with concentrated AuNPs at a molar ratio of 270:1 for 1 hour on ice. Polyethylene glycol (PEG) was added at 6 PEG/nm^3^ to backfilling^12^ AuNPs for 1 hour on ice to stabilize AuNP-BV16. Finally, the modified AuNPs were washed 3 times and characterized by dynamic light scattering (DLS) and UV-vis spectroscopy. For AuNP conjugation with antibody BV11 and LEL, NaHCO_3_ buffer was replaced by 2 mM borate buffer at pH 8.5. The distribution of AuNP modified by anti-JAM-A antibodies and LEL on monolayers was performed by ICC staining to confirm AuNP targeting.

### OptoBBB *in vitro*

For the BBB transient opening *in vitro*, hCMEC/D3 cells were seeded on transwell inserts for about 1 week to form monolayers with a TEER value of around 60 Ω.cm^2^, P_app_ of 8.38±1.17×10^−7^ cm/s for 40 kDa FITC-dextran. The expression of JAM-A on D3 cells was confirmed by ICC staining. The culture was incubated with AuNP-BV16 or AuNP-LEL for 0.5 hours at 37 °C and 5% CO_2_. The treated cellular monolayer was washed three times with PBS before a 28-picosecond laser (532 nm) was applied to the cellular monolayer. The BBB opening *in vitro* was characterized by TEER and permeability measurements.

### WST-1 assay

The hCMEC/D3 cells were seeded in 96-well plates in 100 μL of culture medium for about 1 week to form monolayers. The electron mediator solution and developer reagent were mixed in equal volumes. 10 μL WST-1 mixture was added to monolayers, followed by 2 hours of incubation at 37 °C and 5% CO_2_. After incubation, the 96-well plates were wrapped up in foil and shaken for 1 minute gently at room temperature. Finally, the absorbance of samples was detected at a wavelength of 450 nm with a microplate reader (Synergy2, BioTek).

### AuNP biodistribution *in vivo*

Briefly, the mice were intravenously administered AuNP-BV11 or AuNP-LEL, or the control group AuNP-PEG, with a dose of 18.5 μg/g. At 1 hour, we performed cardiac perfusion PBS and collected the brain tissue. The tissue was then digested in fresh-made aqua regia and centrifuged at 5,000 g for 5 minutes to collect the supernatant. Then the gold concentration was measured by Inductively Coupled Plasma Mass Spectrometry (ICP-MS).

### OptoBBB *in vivo*

OptoBBB *in vivo* was performed as described previously^4^. The mice were briefly anesthetized and intravenously administered AuNPs functionalized by BV11 (targeting JAM-A) or by LEL (targeting glycoprotein) with a dose of 18.5 μg/g. Then, the scalp was carefully removed to expose the skull, followed by transcranial picosecond laser stimulation (1 pulse, 28 picosecond pulse duration, 6 mm beam diameter). The molecular tracer 980 Da Evans blue (2% in PBS, 100 μL) was injected intravenously 5 minutes before the laser. After 0.5 hours, transcardial perfusion was performed with 25 mL PBS and then 4% PFA. The brains were extracted to visualize the extravasation of Evans blue.

### Ca^2+^ signal detection

The fluorescent change of fluo-4 represents the cytosolic Ca^2+^ concentration change. The fluorescent intensity of fluo-4 was detected by an inverted microscope (IX 73, Olympus) combined with a fluorescent illumination system and HCImage Live software. The illumination is from X-Cite 110LED Illumination System (Lumen Dynamics Group Inc.). Fluo-4 was excited at 475 nm, and the emission was collected at 509 nm. All the videos were recorded with the same illumination conditions. The images were captured at 1 frame/second with a digital camera (Hamamatsu ORCA-Flash4.0 LT C11440). For Ca^2+^ signal detection, D3 cells were first seeded on transwell inserts to form monolayers as described above. Monolayers were then incubated with AuNP-BV16 or AuNP-LEL at desired concentration for 0.5 hours and then with 3 μM fluo-4 indicators after washing away unbound AuNPs. The inserts were then cut off and placed above the 35-mm dishes. Next, the picosecond laser with a focused beam was aligned with the 10X objective. Lastly, the intensity of fluo-4 was read before and after laser excitation, while the camera was blocked from the laser during laser excitation.

### Ca^2+^ data analysis

The Ca^2+^ imaging data were analyzed using custom Python (https://www.python.org/) code implemented inside a Jupyter notebook^46^. Images were processed and analyzed using the scikit-image library^47^. The analysis also made use of the NumPy and pandas libraries^48, 49^.

First, regions of interest corresponding to individual cells were identified by an iterative image segmentation process. Starting with the image three frames before the laser was applied, every other consecutive image in the data set was segmented to identify a set of cell regions of interest at that time point corresponding to increases in the fluorescence intensity over the background noise and the resting fluorescence levels before application of the laser. The segmentation procedure consisted of identifying local maxima in the fluorescence intensity after subtracting the resting fluorescence and computing an elevation map using a Sobel filter; included local maxima were required to be at least 20 pixels (12.8 μm) from other maxima. The local maxima were used as markers along with the edges detected by the Sobel filter to segment the image using a watershed algorithm. Then distinct regions were filled and labeled to identify the cell regions of interest. Small regions with an area less than 50 pixels were then pruned from the set to avoid including regions significantly smaller than the expected cell size. At each time point, the identified regions were merged with the set of regions identified at the previous time point to iteratively update the set of regions for cells as they are activated over time. When merging the sets of regions, any two regions with a centroid less than 20 pixels (12.8 μm) from one another were combined into a single region, assuming they are highly overlapped and thus correspond to the same cell. The final set of regions was then further filtered based on several region properties to remove any spurious regions, including those with areas less than 100 pixels or greater than 750 pixels, major or minor axis widths less than 8 pixels, eccentricity greater than 0.9, or extent less 0.5 were removed.

Next, the change of fluorescence (ΔF/F) of fluo-4 for each identified cell region was computed by the change in fluorescence, Δ*F* (the intensity at a time t, *Ft*, minus the baseline intensity, *F*), divided by the baseline intensity *F*. The image domain was then divided radially into bins of width 50 μm with origin at the center of the laser spot, and the ΔF/F trace of cells within each radial bin was averaged to estimate the Ca^2+^ signal as a function of distance from the center of the laser spot.

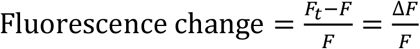

### Western blot

To study the ERK1/2 phosphorylation (p-ERK1/2), we incubated monolayers with targeting AuNPs for 0.5 hours, followed by laser stimulation. We then performed total protein extraction at various time points. For the blocker U0126 (inhibitor of p-ERK1/2) application, the monolayers were pretreated with 10 μM U0126 for 1 hour before AuNP incubation. In the absence of AuNP incubation, proteins were collected at 0.5 hours after laser stimulation as only laser control. We followed the manufacturer’s instructions to extract proteins. Briefly, total proteins from hCMEC/D3 cells were extracted using RIPA buffer (89900, Fisher Scientific) following the manufacturer’s instructions. D3 cells were washed twice with cold PBS, followed by incubation of cold RIPA buffer mixed with a protease inhibitor cocktail (78410, Fisher Scientific) and a phosphatase inhibitor cocktail (78420, Fisher Scientific) for 5 minutes on ice. The lysate was centrifuged to collect the supernatant. Total protein concentration was determined by the BCA assay kit (23225, Fisher Scientific). The exact amounts of proteins (10 μg) were loaded to stain-free protein gel (5678093, Bio-Rad) and separated with electrophoresis under 110 V. The proteins were then transferred to LF PVDF membrane (1704275, Bio-Rad) with the Trans-Blot Turbo system (Bio-Rad), followed by blocking with 5% BSA in TBS buffer at room temperature. After 1 hour, the membranes were incubated with primary antibodies at 4 °C overnight. After washing the primary antibodies with TTBS buffer, corresponding secondary antibodies were applied at room temperature for 1 hour. The blots were enhanced with chemiluminescence (1705060, Bio-Rad), then visualized with ChemiDoc Touch Imaging System (Bio-Rad). The immunoblots were analyzed using ImageLab (Bio-Rad).

## Supporting information

Supplementary Materials for Calcium-Mediated Modulation of Blood-Brain Barrier Permeability by Laser Stimulation of Endothelial-Targeted Nanoparticles

Supplementary Video S1 for Calcium-Mediated Modulation of Blood-Brain Barrier Permeability by Laser Stimulation of Endothelial-Targeted Nanoparticles

## Author Contributions

Author Contributions: Xiaoqing Li designed and performed the experiments for this work and wrote the manuscript. Qi Cai participated in the cell culture, WST-1, and calcium detection. Blake Wilson participated in the calcium analysis. Hanwen Fan participated in the gold nanoparticle conjugation. Monica Giannotta developed the anti-JAM-A antibodies (BV16 and BV11) for this study. Monica Giannotta, Robert Bachoo, and Zhenpeng Qin supervised the project. All authors provided critical feedback and helped shape the research, analysis, and manuscript.

## Competing interests

A patent has been filed related to the technology described in this work (US 2021/0252151 A1).

## Acknowledgments

The authors thank Yaning Liu for assistance with picosecond-laser alignment and members of the Qin laboratory for discussions, Dr. Theodore J. Price and Ayesha Ahmad for assistance with western blot. This research was funded by Cancer Prevention and Research Institute of Texas (CPRIT) grants RP160770 and RP190278, American Heart Association Collaborative Sciences Award (19CSLOI34770004), European Research Council (project EC-ERC-VEPC, contract 742922) and a Fondazione CARIPLO Foundation grant (2016-0461).

