## Supplementary Materials for Calcium-Mediated Modulation of Blood-Brain Barrier Permeability by Laser Stimulation of Endothelial-Targeted Nanoparticles for "Calcium-Mediated Modulation of Blood-Brain Barrier Permeability by Laser Stimulation of Endothelial-Targeted Nanoparticles"

[1]Department of Bioengineering, the University of Texas at Dallas, Richardson, TX 75080, USA
[2]Department of Mechanical Engineering, the University of Texas at Dallas, Richardson, TX 75080 USA
[3]Institute of Molecular Oncology Foundation (IFOM), 20139 Milan, Italy
[4]Department of Internal Medicine, the University of Texas Southwestern Medical Center, Dallas, TX75390, USA
[5]Harold C. Simmons Comprehensive Cancer Center, the University of Texas Southwestern Medical Center, Dallas, TX75390, USA.
[6]Department of Neurology, the University of Texas Southwestern Medical Center, Dallas, TX75390, USA
[7]Center for Advanced Pain Studies, the University of Texas at Dallas, Richardson, TX 75080, USA
[8]Department of Surgery, University of Texas Southwestern Medical Center, Dallas, TX75390, USA

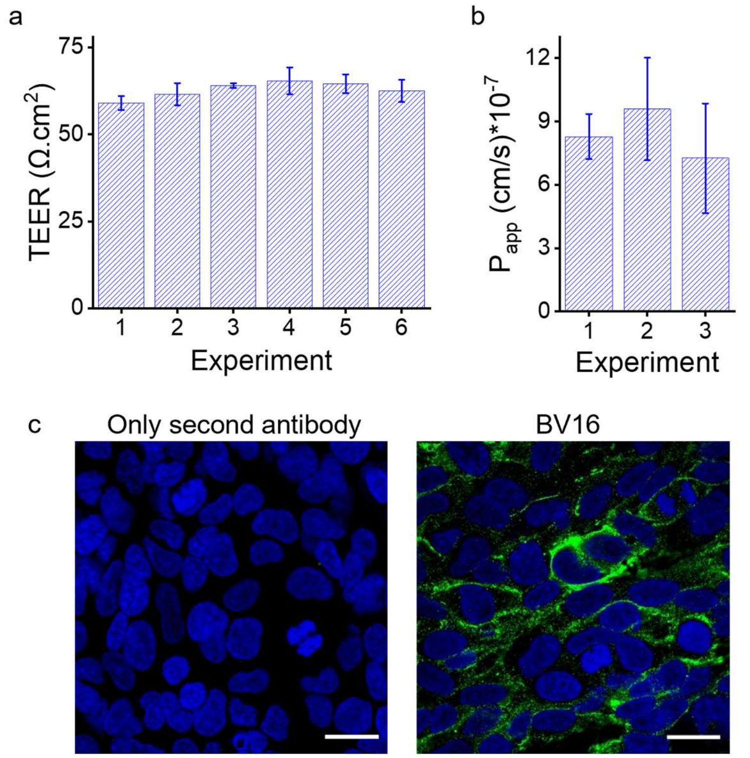


**Figure S1. Characterization of *in vitro* BBB model using hCMEC/D3 monolayers. (a)** Transendothelial electrical resistance (TEER) of D3 monolayers. 3 replicates in each experiment. **(b)** Permeability to FITC-dextran (40 kDa) of D3 monolayers. 3 replicates in each experiment. **(c)** Formation of tight junctions detected by ICC staining. Green: JAM-A detected by BV16. Blue: nuclei. Data expressed as Mean ± SD. Scale bar: 10 µm.

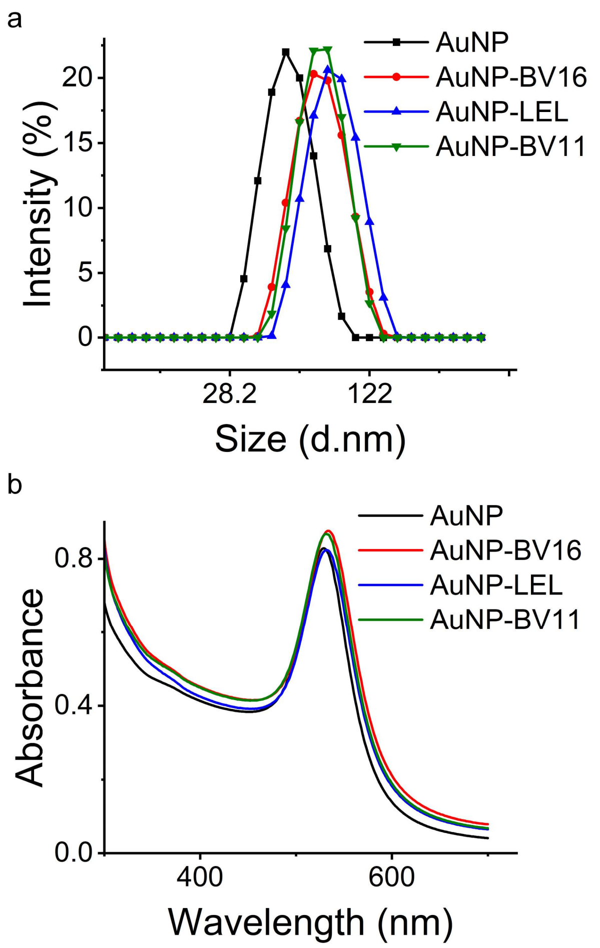


**Figure S2. Characterization of AuNPs modified with different targets.** Characterization of AuNPs modified with BV16 (mouse anti-human JAM-A antibody), LEL (*Lycopersicon esculentum* lectin for targeting glycoprotein), and BV11(rat anti-mouse JAM-A antibody) by DLS (a) and UV-vis spectroscopy (b).

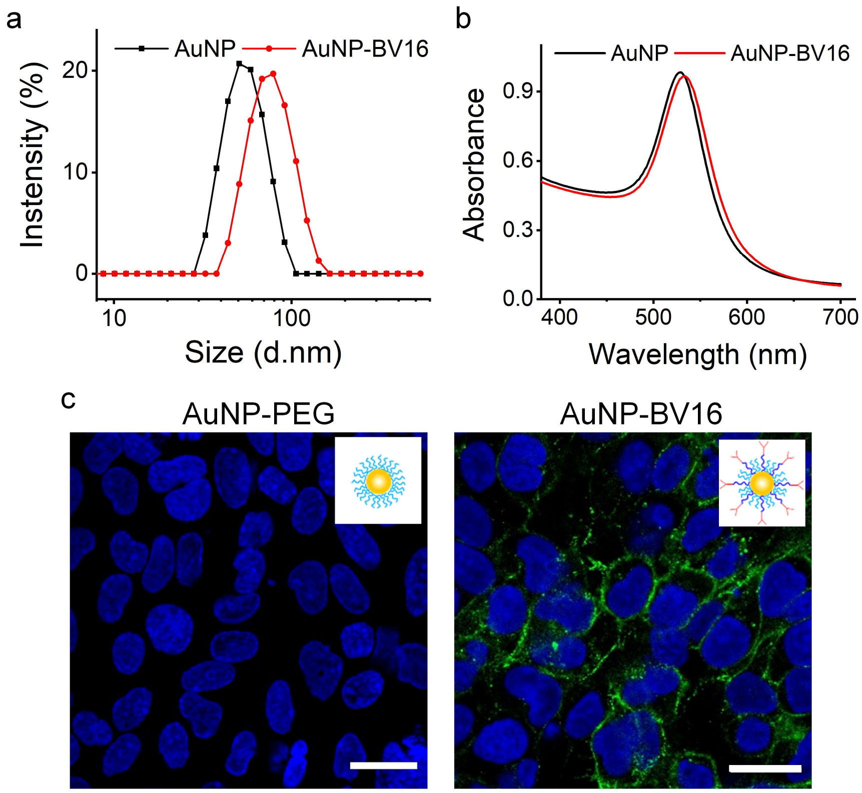


**Figure S3. Characterization and targeting of AuNP-BV16. (a-b)** Characterization of AuNP-BV16 by DLS (a) and UV-vis spectroscopy (b). **(c)** Tight junction targeted by AuNP-BV16 (0.5 nM) using ICC staining, while non-targeting AuNP (AuNP-PEG) does not show specific targeting. Green: JAM-A stained by AuNP-BV16. Blue: nuclei. Scale bar: 10 µm.

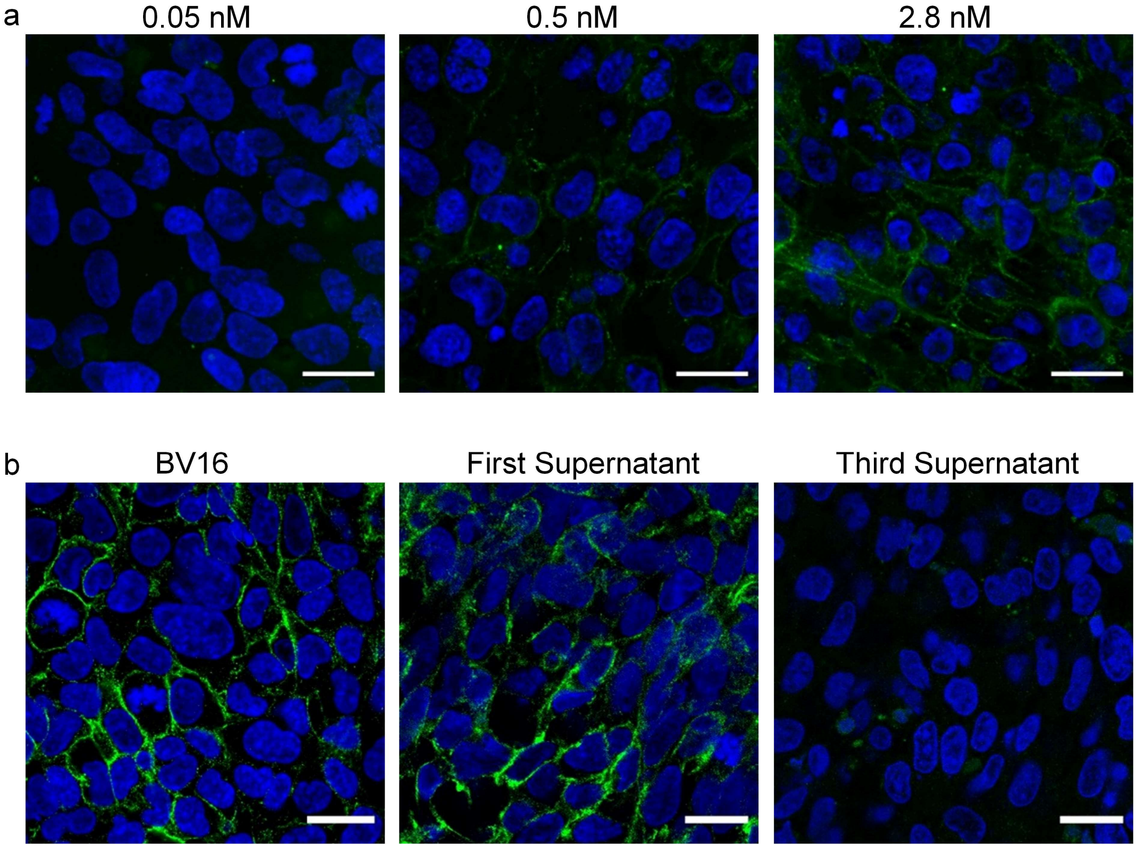


**Figure S4. Optimization of AuNP-BV16 for OptoBBB. (a)** Distribution of TJ-targeting AuNP (AuNP-BV16) on D3 monolayers with various AuNP concentrations. Obvious TJ targeting using 0.5 nM and 2.8 nM AuNP-BV16 was observed, but not 0.05 nM AuNP-BV16. Thus, 0.5 nM AuNP-BV16 was selected for OptoBBB *in vitro*. **(b)** Lack of free antibody in the AuNP-BV16 conjugate. The free BV16 will compete with AuNP-BV16 for JAM-A binding, which affects AuNP-BV16 targeting. The first supernatant and third supernatant were incubated with monolayers, respectively, to confirm if there were free BV16 after washing. The free antibody had been removed after washing 3 times indicated by the right panel image, which doesn't show green signal. BV16: free BV16 antibody as positive control, first supernatant: supernatant from AuNP-BV16 conjugation after washing 1 time. Third supernatant: supernatant from AuNP-BV16 conjugation after washing 3 times. Scale bar: 10 µm.

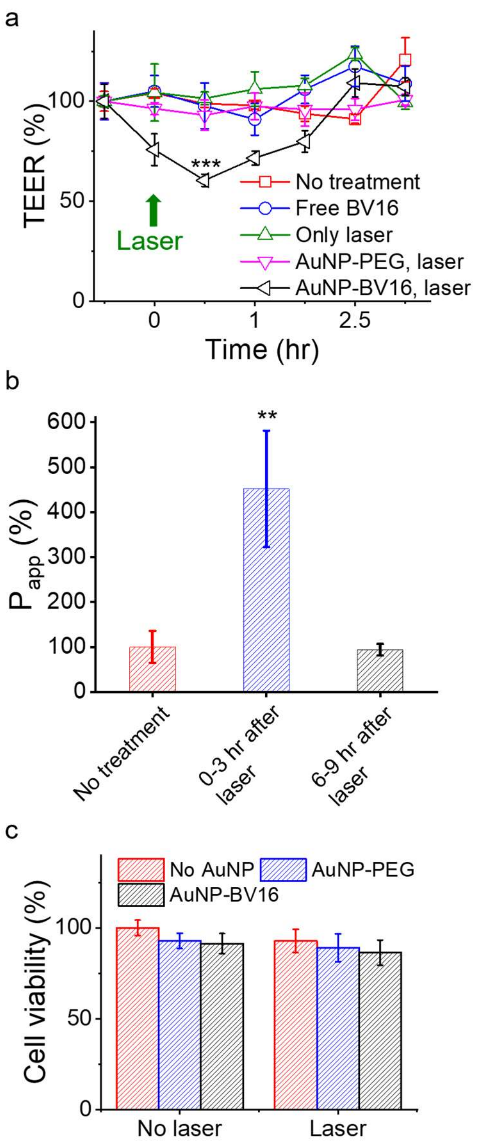


**Figure S5. Reversible BBB opening by laser excitation of AuNP-BV16. (a)** Normalized TEER change over time by laser stimulation. n=3 replicates. **(b)** Reversible permeability changes after laser stimulation. n=3 replicates. **(c)** No significant difference in cell viability after laser stimulation. n=6 replicates. Incubation concentration: 0.5 nM, laser: 35mJ/cm$^2$, 25 pulses, 5 Hz. Data expressed as Mean ± SD. Unpaired *t-test* was performed individually between No treatment (No laser and No AuNP) and the other groups. **: P<0.01, or *** P<0.001was considered a statistically significant difference.

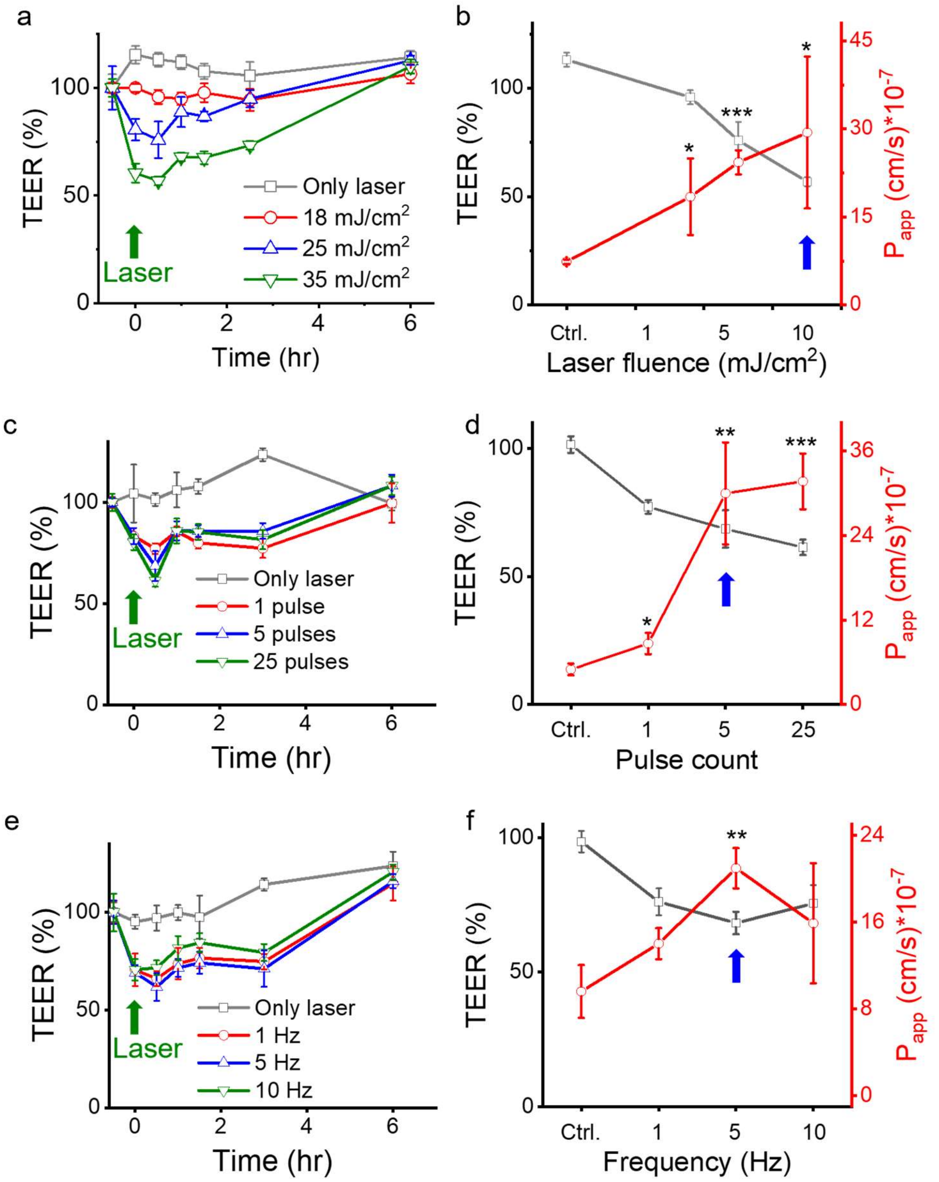


**Figure S6. Effects of laser parameters on OptoBBB. (a-b)** TEER (a) and permeability (b) change after laser stimulation under different laser fluences. 25 pulses, 5 Hz. **(c-d)** TEER (c) and permeability (d) change after laser stimulation under different laser pulse counts. 35 mJ/cm$^2$, 5 Hz. **(e-f)** TEER (e) and permeability (f) change after laser stimulation with different frequencies. 35 mJ/cm$^2$, 25 pulses. Data expressed as Mean ± SD (n=3). Unpaired *t-test* was performed for permeability individually between Only laser and the other groups. **: $P<0.01$ was considered a statistically significant difference. The blue arrow indicates the condition selected for the following experiments.

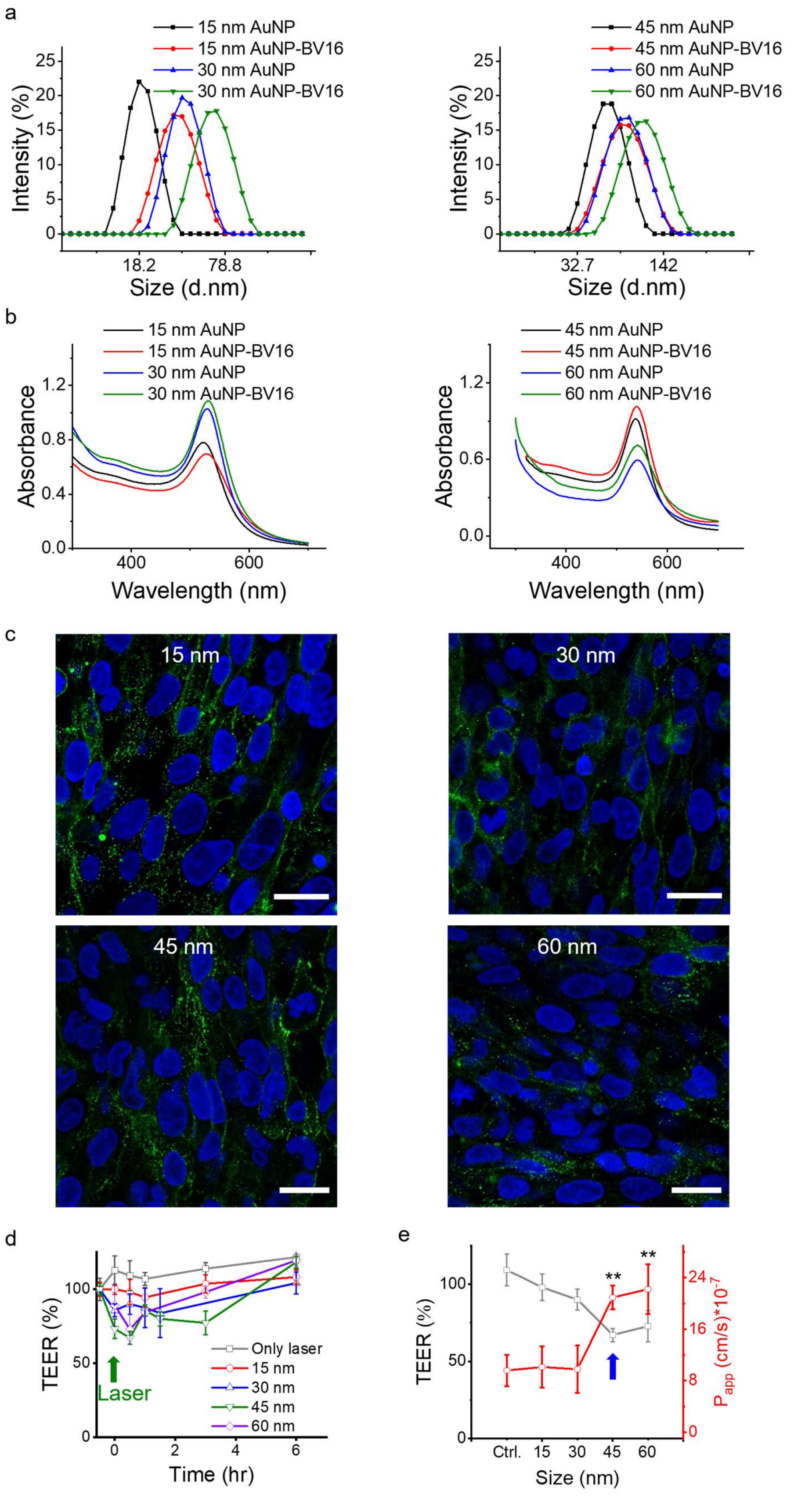
a
15 nm AuNP
15 nm AuNP-BV16
30 nm AuNP
30 nm AuNP-BV16
Intensity (%)
Size (d.nm)
45 nm AuNP
45 nm AuNP-BV16
60 nm AuNP
60 nm AuNP-BV16
b
Absorbance
Wavelength (nm)
c
15 nm
30 nm
45 nm
60 nm
d
TEER (%)
Time (hr)
Laser
Only laser
e
TEER (%)
Size (nm)
Ctrl.
**

**Figure S7. Effects of AuNP size on the BBB opening. (a-b)** Characterization of AuNP-BV16 with particle sizes of 15, 30, 45, and 60 nm using DLS (a) and UV-vis spectrum (b). **(c)** Distribution of AuNP-BV16 on D3 monolayers with sizes 15, 30, 45, and 60 nm. **(d-e)** TEER (d) and permeability (e) change of D3 monolayers with laser excitation of AuNP with sizes 15, 30, 45, and 60 nm. Data expressed as Mean ± SD (n=3). Unpaired *t-test* was performed for permeability individually between Only laser and the other groups. **: P<0.01 was considered a statistically significant difference. The Blue arrow indicates the condition selected for the following experiments. Scale bar: 10 μm.

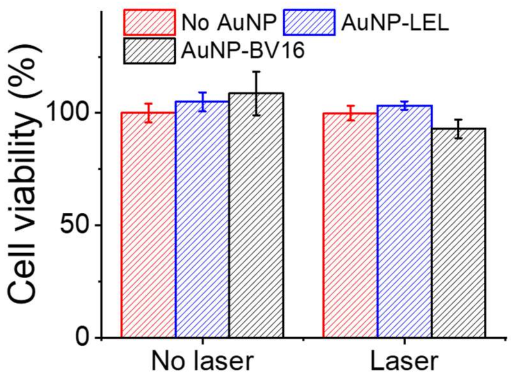


**Figure S8. Cell viability after laser excitation of different targeting AuNPs.** AuNP-LEL: 0.02 nM, AuNP-BV16: 0.5 nM. 35mJ/cm$^2$, 5 pulses, 5 Hz. Data expressed as Mean ± SD (n=6). Unpaired *t-test* was performed for permeability individually between the no treatment (No AuNP w/ No laser) group and the other groups. No significant difference was observed.

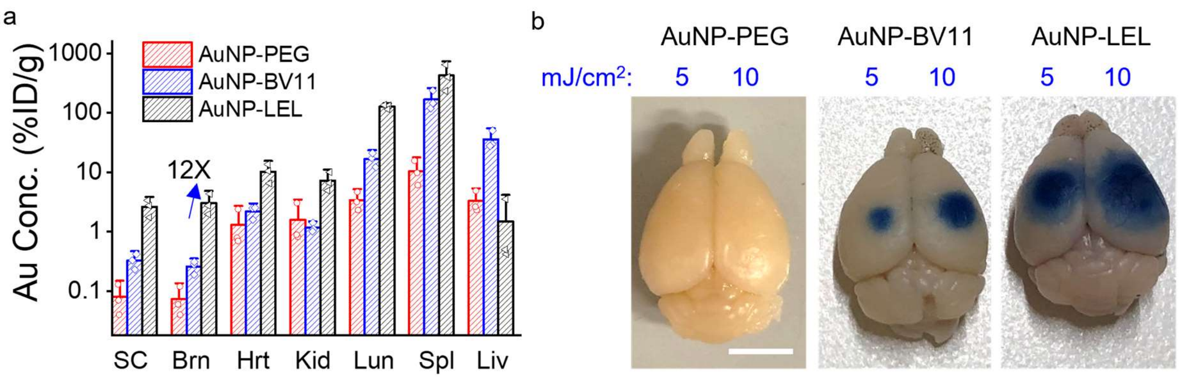


**Figure S9. BBB opening *in vivo* with different targets. (a)** The gold concentration in main organs with different targets. **(b)** Comparison of *in vivo* BBB opening by laser stimulation of AuNP-LEL and AuNP-BV11. 1 pulse. Dose: 18.5 μg/g. SC: spinal cord, Brn: brain, Hrt: heart, Kid: kidney, Lun: lung, Spl: spleen, Liv: liver. Data expressed as Mean ± SD (n=3 mice). Scale bar: 4 mm.

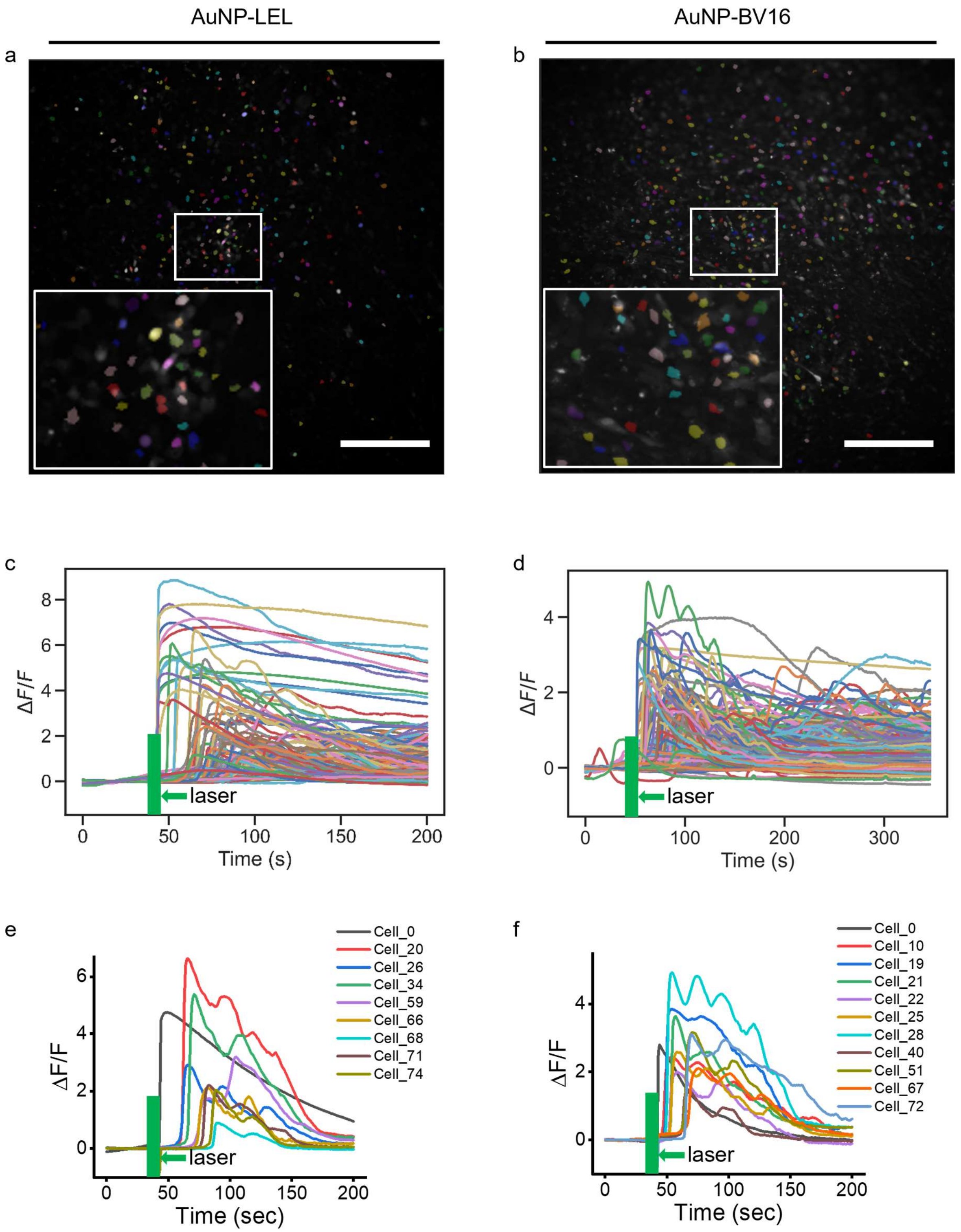


**Figure S10. Representative $Ca^{2+}$ response from one experiment after laser stimulation of AuNP-LEL or AuNP-BV16. (a-b)** Segmentation of samples to show all the cells (labeled by different colors) that involve cytosolic $Ca^{2+}$ elevation after laser stimulation. **(c-d)** The quantification of the $Ca^{2+}$ signal of cells in (a) or (b). **(e-f)** Oscillatory increase in $Ca^{2+}$ of cells in (a) or (b). 35mJ/cm$^2$, 1 pulse. Scale bar: 0.5 mm.

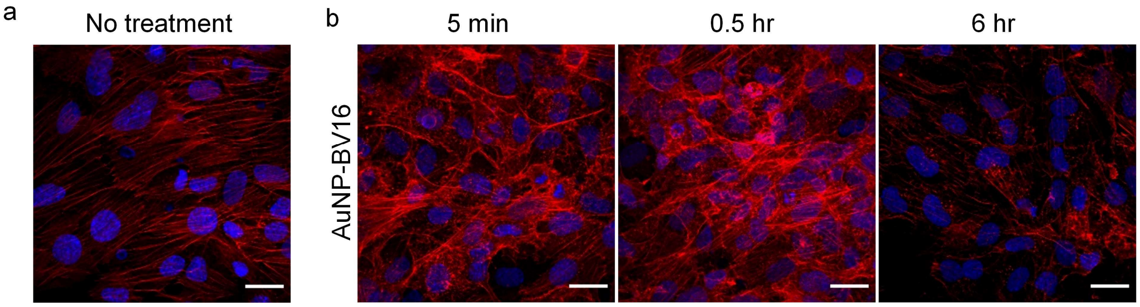


**Figure S11. Actin polymerization after laser stimulation of AuNP-BV16. (a)** F-actin staining of no treatment as control. **(b)** F-actin staining at 5 min, 0.5 hr, and 6 hr after laser stimulation of AuNP-BV16. 35 mJ/cm$^2$, 1 pulse. Red: F-actin stained by phalloidin. Blue: nuclei. Scale bar, 20 μm.
